# Quantius: Generic, high-fidelity human annotation of scientific images at 10^5^-clicks-per-hour

**DOI:** 10.1101/164087

**Authors:** Alex J. Hughes, Joseph D. Mornin, Sujoy K. Biswas, David P. Bauer, Simone Bianco, Zev J. Gartner

**Author notes:** [* Co-first author].

## Abstract

We describe Quantius, a crowd-based image annotation platform that provides an accurate alternative to task-specific computational algorithms for difficult image analysis problems. We use Quantius to quantify a variety of computationally challenging medium-throughput tasks with ~50x and 30x savings in analysis time and cost respectively, relative to a single expert annotator. We show equivalent deep learning performance for Quantius- and expert-derived annotations, bridging towards scalable integration with tailored machine-learning algorithms.

---

Image analysis is increasingly crucial to quantitative biology^1-3^, personalized medicine^4,5^, and disease diagnostics^6,7^. Many image features can be accurately annotated by individual researchers, but this approach becomes impractical for image sets that are readily acquired in the thousands per experiment. Researchers thus resort to custom-written scripts to analyze image features of interest using methods such as contrast-based segmentation, edge detection, and tracking^3,8,9^. These approaches can give excellent performance for highly consistent imaging conditions, but are rarely generic enough to be portable to experiments with slight variations in setup (including different cell lines, fluorescent labels, or imaging magnification), much less completely different image analysis problems. On the other hand, machine learning approaches can be more flexible to different image inputs, but the convolutional neural networks that are currently favored for modern object detection efforts demand extensive datasets of human-annotated images for training purposes^6,7,10^.

The scientific community has begun to build annotation pipelines that leverage the accuracy and throughput benefits of large groups of human annotators working in parallel^11^. Platforms such as Zooniverse^12^, EyeWire^1^, and Foldit^13^ are usually custom-built for specific large-scale projects, and often rely upon volunteer labor^14^, making them cumbersome to implement generically or on small-to-medium scale projects. Other human interaction services exist that could circumvent these problems, such as Amazon’s “Mechanical Turk”. However, these services are not currently used extensively by the life sciences community, even though the use of payment appears to address a variety of problems associated with volunteer labor, including worker dropout, temporally volatile overall contributions, and a tendency for workers to contribute to new, well publicized, or “game-ified” projects in preference to others^15-18^. Additionally, the reliability of such annotations for scientific problems in the life sciences is unclear.

Many projects in the basic sciences rely on iterative cycles of imaging and quantitative analysis, where the objectives of analysis are refined over time in tandem with the underlying experiment design. In these cases, algorithm development is a bottleneck to discovery. A clear path to obtaining large-scale, reliable, high-quality annotation of images for life science projects is currently missing. To this end, we developed Quantius, a flexible portal that enables scientists to leverage groups of untrained mechanical <u>turk</u> work<u>ers</u> (“turkers”) using a set of interaction tools that can be applied individually to many job types. Moreover, annotations can be collected and used in series to refine further rounds of annotation^19^, as pre-segmented input to conventional algorithms, or to train machine learning algorithms (**Fig. 1a**). We created a publicly available website, Quanti.us (*login details will be available on publication, see* ***Methods***), that allows researchers to upload image sets, choose an analysis tool, and provide simple sets of instructions with example images to turkers. Images can be presented to turkers as individual images or image stacks. Each image or stack is referred to as a “human intelligence task” (or simply “task”) within a larger “job”. The website acts as a generic portal, automatically interfacing with mechanical turk to setup tasks and return data relating to the clicks that turkers made in each task. These data fields include click Cartesian coordinates, anonymized turker identification numbers, and time-stamps.

**Figure 1.**
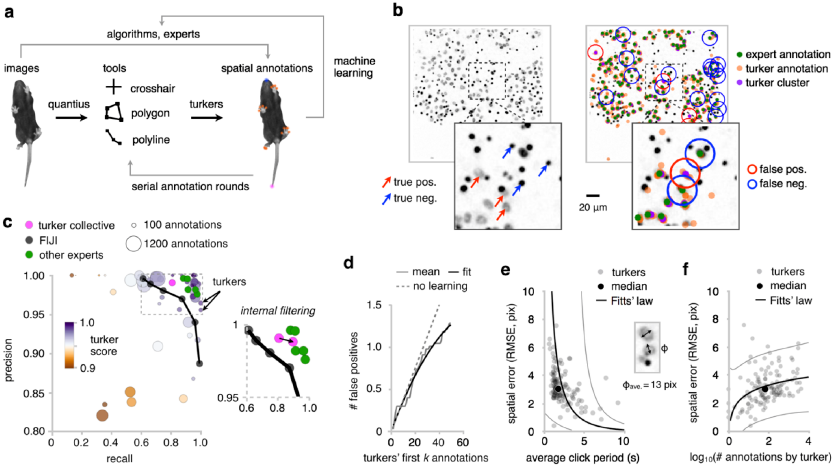
Accurate hand annotations that leverage the “wisdom of crowds” for nimble and generic scientific image analysis using Quantius. **(a)** Scientists choose an interaction tool that human ‘turkers” use to annotate uploaded images according to a brief instruction set. These annotations Dan be interpreted in raw form, passed as input to conventional algorithms, or as input to machine earning routines such as convolutional neural networks, **(b)** Turkers were tasked with discriminating oetween cell nuclei (true positives) and autofluorescent pores (true negatives) using a crosshair tool; false positive and negative annotations were scored against a trained expert for individual turkers or for spatially clustered^20^ annotations from all turkers. Each of 300 images was annotated by 10 turkers (a subset of 20 was used to extract turker performance metrics), **(c)** Precision and recall, metrics for individual turkers, for clustered annotations (“turker collective”), for other experts, and for a conventional FIJI object detection pipeline over a range of particle size thresholds (n = 46 turkers). Turkers could be assigned an inherent quality score relative to their peers, independent of the expert dataset. Inset arrow: effect of filtering out the bottom third-performing workers, **(d)** False oositive errors contributed over the first *k* annotations a turker submitted (in chronological order) fit oy a quadratic function, indicative of a linearly decreasing error rate (*n* = 29 turkers). **(e)** Spatial error af annotations shows a Fitts’ law tradeoff with the time between annotations (*n* = 129 turkers). **(f)** Average spatial errors increase weakly with the number of annotations a turker contributes, but are small relative to the average nuclear diameter. Fit envelopes in **(e)** and (f) are 95% confidence ntervals.

We first evaluated Quantius for a challenging particle discrimination task, in which fluorescently-tagged cells migrate through a porous membrane in a transwell assay (**Fig. 1b**). Counting cells using conventional contrast-based segmentation in e.g. the National Institutes of Health’s FIJI is difficult because the pores of the membrane are autofluorescent, making them hard to distinguish from cells. We tasked turkers with clicking on cells using a crosshair tool, and compared their performance to traditional FIJI segmentation and a “gold-standard” expert dataset. We found that 27 of 46 turkers (59%) that completed at least one image performed better in both precision (measuring the ability of turkers to exclude false positives) and recall (measuring their ability to exclude false negatives) than a custom FIJI pipeline consisting of brightness threshold, watershed, and particle size threshold steps (**Fig. 1c**). Moreover, the use of multiple turkers to analyze any given image provides significant additional advantage at reasonable cost. By using subtractive spatial clustering^20^ of the annotations from the 10 replicate turkers that were tasked with analyzing each image, we leveraged the “wisdom of crowds”^17^ to produce precision and recall metrics of 0.99 and 0.81; 0.015 and 0.18 higher, respectively, than the performance envelope followed by the FIJI algorithm for different particle size thresholds. We found that the advantage of crowds was due to diluting the impact of rare, poor-performing workers, and was accrued for group sizes of as little as 3 (**Supplementary Fig. 1**). We also generated an internal turker quality score by comparing annotations from each turker with clusters generated from those made by their peers, which can enable poorer-performing turkers to be automatically screened out, even without an expert dataset to compare to^21^. Filtering out the contributions of the bottom third of the turkers based on this internal quality score made the performance of the turker collective indistinguishable from that of a group of 5 other experts (**Fig. 1c, inset**).

Consistent with other studies of worker contributions in crowds, distributions describing the number of images attempted by turkers generally followed the Pareto “80/20” principle, namely that ~20% of turkers accounted for ~80% of images completed^11,15^ (**Supplementary Fig. 2**). Turkers appeared to improve in performance dynamically at the outset of the cell/pore discrimination task, reducing their average false positive rate approximately linearly from 3.7% to 1.3% per annotation in the first 50 annotations that they made (**Fig. 1d**). A linear reduction in false positive rate with training time is consistent with other types of object recognition tasks in humans^22^. These data suggest additional strategies for collating turker data for improved accuracy, such as by providing an initial training image set, or by weighting later annotations in a particular job more heavily than early annotations.

**Figure 2.**
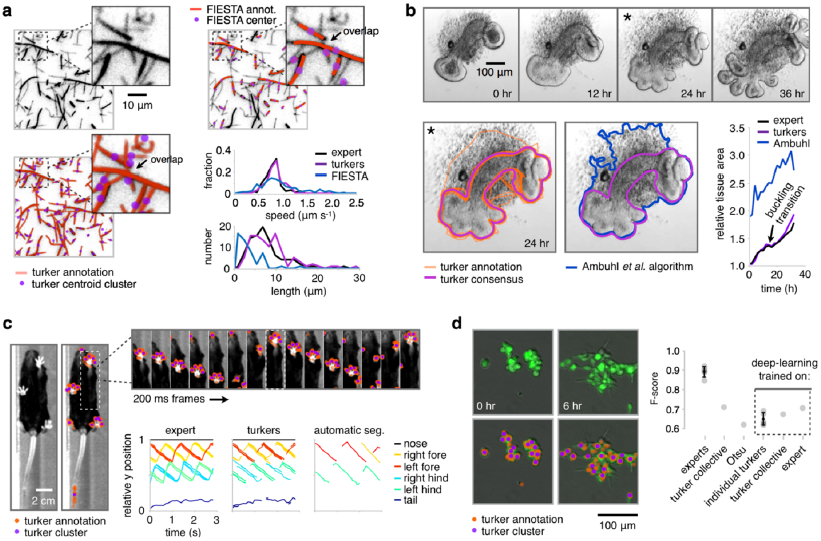
Performance and machine-learning integration of Quantius for image features that overlap or are difficult to segment, **(a)** Turkers were tasked with tracing fluorescent microtubules in a “gliding” assay using a polyline tool. Each of 50 images was annotated by 10 turkers. Polyline centroids were spatially clustered and used as input to FIJI’S TrackMate to determine velocity distributions relative to hand annotations by a trained expert, or to FIESTA output. Length distributions were determined by Deming regression of clustered polylines^33^, **(b)** Turkers were tasked with outlining the smooth epithelial layer of a lung explant (images reproduced from ref. ^27^). Each of 17 images was annotated by 20 turkers. A consensus outline was determined after spatial clustering of outline centroids using a sliding spline fit. The consensus outline matches tissue area estimates from outlines collected by a trained expert, relative to a conventional segmentation algorithm from Ambuhl *etal.*^29^. **(c)** Turkers were tasked with annotating the nose, digits, and tail of a live behaving mouse (images reproduced from ref. ^34^). Each of 29 images was annotated by 20 turkers. Annotations from a trained expert or spatially clustered turker annotations were passed as input to FIJI’S TrackMate to construct a gait plot tracking anatomical feature positions overtime. An incomplete plot was recovered through a conventional segmentation pipeline in FIJI consisting of brightness thresholding, particle analysis, and TrackMate. **(d)** Clustered turker annotations were used to train a convolutional neural network to surpass the performance of conventional segmentation for 48 frames from a movie of mammary epithelial organoid spreading *in vitro* (10 turkers per frame). A neural network trained on annotations from a 10?turker collective gave similar performance to those trained on expert annotations, and all networks performed better than traditional Otsu segmentation.

Turkers chose to annotate images with different average click periods, which correlated with their spatial root-mean-square errors (RMSEs). Spatial RMSE is a measure of click accuracy, here evaluating the average spatial distance between the clicks that a turker made, and the corresponding clicks by an expert. Median turker click times were ~2 s for RMSE’s of ~3 pix (**Fig. 1e**). The scaling between the two showed that turkers that spent more time on each click had lower (more accurate) RMSE’s, following Fitts’ law of speed-accuracy tradeoffs in human pointing tasks^23,24^. Secondly, turkers that contributed more annotations to a job tended to choose smaller click periods, leading to increased (less accurate) RMSE’s (**Fig. 1f**), although other factors such as fatigue may have contributed to this relationship. However, all 129 turkers that made at least one annotation had RMSE’s below 13 pixels (the average diameter of the cells they annotated), indicating high overall pointing performance.

Because Quantius pays individuals to annotate images, turkers make decisions based largely upon the economic tradeoffs inherent in completing a task accurately and quickly. We therefore explored the relationships between turker performance, task complexity, overall task completion rate, and the amount turkers were paid per task. Here we ran a separate set of calibration experiments on synthetic images containing different numbers of particles distributed spatially according to a known ground truth. We found that turkers tolerated around 60 annotations per image task at a pay of 2¢ per image (USD, **Supplementary Fig. 3a-d**). Above this threshold in image complexity, the average number of images attempted by turkers dropped from 15 to 8, while recall dipped from >0.8 to ~0.6 above 110 particles per image. The total rate at which annotations were collected dropped from >25,000 hr^-1^ to ~3,000 hr^-1^ across the same threshold. However, increasing the pay per image broadly reversed these losses in recall and overall annotation rate incurred by increased image complexity (**Supplementary Fig. 3e-g**). Turkers tended to favor higher click speeds (and higher spatial error) for more complex images or lower pay per image, but each metric only varied within a ~20% envelope across the wide range in complexity and pay parameters we studied.

Notably, we achieved an annotation collection rate of ~105 hr^-1^ for images requiring 110 annotations each at 6¢ per image (**Supplementary Fig. 3e**). These metrics compare favorably to those of an individual academic or industry researcher, who would need to spend $800-$2,000 USD in salary and ~56 hours to make the same number of annotations collected for $55 in 1 hr using Quantius (~50x and 30x savings in analysis time and cost, respectively, **Supplementary Note 1**).

Beyond object discrimination tasks that require independent clicks using a crosshair, we challenged turkers with more complex job types that required multiple crosshair, polyline, or polygon annotations. First, we asked turkers to draw polylines over fluorescently labeled microtubules recorded in a “gliding” motility assay^25^ (**Fig. 2a**). Here, microtubules move over the surface of a glass slide coated with dynein motors, and present a significant challenge to automated segmentation efforts because of the similarity in brightness between microtubules that often overlap. We spatially clustered polyline annotations from 10 turkers per image, and used these cluster centers as input to FIJI’s TrackMate plugin^26^. In an independent analysis, we applied an algorithm custom-written for quantification of microtubule gliding assays known as the FIESTA^9,25^. Although both Quantius and FIESTA velocity distributions approximately matched that recovered from manual microtubule tracking by a human expert, the Quantius microtubule length distribution matched the expert distribution more closely than FIESTA’s. This appeared to be because turkers were more adept at successfully ignoring overlap junctions between different microtubules, while FIESTA tended to break microtubule annotations into smaller segments bordered by junctions.

Second, we asked turkers to draw closed polygons over frames from a phase-contrast movie of an epithelial lung explant undergoing branching morphogenesis in vitro^27^ (**Fig. 2b**). Although the development of algorithms applicable to drawing an arbitrary boundary through outlines aggregated across different workers is an open area of research^28^, we used a centroid clustering approach to reject errant outlines. This approach again leveraged the “wisdom of the crowd” to generate consensus outlines suitable for comparison against outlines from a conventional phase-contrast segmentation algorithm^29^, or from a trained expert. The consensus turker outlines and the expert outlines gave similar estimates of the tissue area over the course of the movie, and successfully captured a stall in morphogenesis just prior to a mechanical buckling transition in the tissue. On the other hand, the automatic algorithm could not distinguish the outline of the epithelium from background in the images caused by migratory cells in adjoining tissue layers, leading to noisy and inaccurate area estimates.

Third, we asked turkers to make multiple crosshair annotations to track the nose, digits, and tail of a freely moving mouse in a movie of its ventral aspect (**Fig. 2c**). These annotations were spatially clustered and used as input to FIJI’s TrackMate, successfully capturing much of the dynamics recovered through manual gait analysis by a trained expert. On the other hand, conventional contrast-based segmentation and tracking methods in FIJI were hindered by the high variation in contrast between the anatomical features of the mouse and the background over the length of the movie. Here, only two of the paws could be tracked consistently, and not with single-digit resolution.

Beyond the use of Quantius as a tool for rapidly generating large sets of human annotations for diverse scientific image types, we explored its integration with machine learning. The integration of Quantius with a generic machine learning pipeline could provide an economical alternative to hand annotation of very large image sets^15^, since hand annotations could be collected only until sufficient to train a machine with acceptable performance for subsequent stand-alone tasks of the same type. As a proof of principle of this capability, we studied a movie of fluorescently-labeled mammary epithelial cell clusters spreading over an *in vitro* culture surface (**Fig. 2d**). We expected this to be a particularly challenging computational problem because of the wide range in cell morphologies observed over the course of the movie, and the spatial overlap between cells in the clustered state.

We trained deep convolutional regression networks^30-32^ with turker annotations to determine whether they could generate algorithms with comparable performance to those trained on annotations from trained experts. Indeed, an algorithm trained on the annotations of 10 turkers gave a comparable F-score (the harmonic mean of precision and recall) to an algorithm trained on the annotations of an expert, at 0.68 and 0.71 respectively. Both algorithms showed better performance than traditional Bayes-optimal Otsu segmentation, a non-machine learning approach (F-score of 0.62). Further, the algorithm trained on the turker collective performed better than the mean performance of algorithms trained on annotations from individual turkers, reflecting a “wisdom of the crowd” benefit to the training process that was mainly accounted for by increases in recall (**Supplementary Fig. 4**).

The current demand for quantitation in the life sciences necessitates nimble and integrated approaches to image analysis and assay development; especially for life science researchers that are not themselves experts in image analysis, or for problems that remain challenging to address using computational algorithms. Moreover, the increasing practicality and availability of supervised machine learning creates a significant opportunity to dispense with continuous development of custom-written object detection algorithms. We found that hand annotations could be collected at significant scale and quality using Quantius, marshaling paid turkers to label any of a range of image sets using a core set of interaction tools. These annotations are invaluable for difficult segmentation tasks, both for rapid pilot-scale analyses, and towards relieving dataset volume bottlenecks in tailored convolutional neural networks. The Quantius platform can be further improved to meet the needs of the biomedical research community. For example, additional annotation tools could be added. We believe that even higher-quality pools of turkers could be curated by dynamically tracking turker performance using intrinsic metrics. Further, Quantius tasks could be daisy-chained and integrated with machine learning to produce multi-stage annotation pipelines. These efforts would significantly simplify, and thereby democratize, quantitative biology analyses for fundamental, health-related, and diagnostic ends.

## Methods

### Annotation collection using Quantius

Image sets were uploaded to <u>quanti.us</u>, a publicly available website developed in this work (*currently in a beta testing stage during manuscript review*). The website enables a user to upload an image set (each image is a *task* within the larger *job*), select an annotation tool, provide instructions to turkers (**Supplementary Fig. 5**), and specify the desired number of *replicates* (number of independent turkers making annotations on each image task). Each image task in the job is created as a "human intelligence task" on Amazon mechanical turk^35^. When all tasks are complete, Quantius returns annotations as Cartesian coordinates, along with individual anonymized turker identification numbers. Time-stamps are also returned for each submitted image, and for each annotation relative to the first annotation made by a given turker on that image.

### Annotation post-processing

Annotations were overlaid onto images and post-processed using custom scripts in MATLAB R2015b (Mathworks, Natick, MA). Annotations by different turkers for the same image were spatially grouped by subtractive clustering (subclust.m). Clustering was performed on individual annotations made with the crosshair tool, or on centroids of sets of annotations made with the polyline or polygon tools. When analyzing crosshair annotations by individual turkers, false positive annotations (*fp*) were taken to be those that fell further than *x* pixels from the nearest annotation in the corresponding expert “gold-standard” dataset, while false negatives (*fn*) were taken to be gold-standard annotations that fell further than *x* pixels from the nearest annotation made by the turker. The value of *x* was set to twice the approximate average full-width-at-half-maximum of the objects being annotated. Collective performance of turkers was evaluated using similar computations for clusters rather than for individual annotations. A simple turker score was defined as 1 - [(*fp*’ + *k*.*fn*’)/(2.*tp*’)], where *fp*’ and *fn*’ are false positives and false negatives determined for the turker under consideration relative to turker annotation clusters rather than the expert gold-standard annotations. The score can be tailored to specific job types through the arbitrary parameter *k* (we set *k* = 0.2 for **Fig. 1**). More complex inherent worker quality scores have also been defined^21^.

Spatial root-mean-square errors (RMSEs) for each turker were computed from the distances between their true positive annotations and corresponding gold-standard annotations. Performance metrics were precision = *tp*/(*tp*+*fp*) and recall = *tp*/(*tp*+*fn*). Fitts’ law was fit using the Levenberg-Marquardt algorithm^36^ (fit.m) as *t* = *a* + *b*.log_2_[(*c* +σ)/ σ], where *a, b*, and *c* are fitting parameters; and σ is taken to be RMSE. For Fig. 1f, we assume t ∝ -log(*# annotations done*), see (**Supplementary Fig. 6**), such that Fitts’ law can be trivially recast as: log(*# annotations done)* = *d* - *e*.log_2_[(*c* +σ)/ σ]; where *d, e*, and *c* are fitting parameters.

The centroids of polyline and polygon objects associated with each image were clustered and an arbitrary distance threshold used to determine membership of a given polyline/polygon into a group describing a putative image structure. Polylines/polygons outside this threshold were discarded. For microtubule polylines, centroids were computed from the set of annotations in each group and passed as input to FIJI’s TrackMate^26^ (National Institutes of Health, Bethesda, MD) to determine velocity distributions. The annotations in each group could also be fit using Deming regression^33^ to extract length distributions (deming.m). For polygon outlines, a consensus outline was determined by first taking an arbitrary outline in the group, and for each annotation in that outline, generating a local centroid from nearby points in other outlines (within an arbitrary distance threshold). These local centroids were then fit using a periodic interpolating cubic spline (cscvn.m), which constituted the consensus outline.

### Microtubule gliding assay

Dynein was purified from squid optic lobes, as previously described for kinesin^37^, except that AMP-PNP was not used during microtubule sedimentation. Optic lobes were dissected and homogenized using a Dounce tissue grinder. The homogenate was pelleted and microtubules and motor proteins extracted from the supernatant via sucrose gradient centrifugation. Taxol-stabilized microtubules were labeled with rhodamine and imaged in dynein-adsorbed glass flow cells by widefield fluorescence microscopy in the presence of ATP.

### Machine learning

We trained a machine learning system to predict the center locations of cell nuclei in 500 ×500-pixel test image. Conventional approaches like extracting regional features (for example, *extremal region*^38^) to develop a region-level-detector did not offer viable options owing to the small size of the cellular objects. In contrast, a convolutional neural network (CNN) allowed an end-to-end system design without extracting separate features to feed into the learning system. The CNN transforms the image channels by applying a set of “learnable” filters, successively, directly into an output matrix (as big as the input image) containing high scores in the locations of nuclei centers and very small to almost zero scores elsewhere. The objective of a fully-convolutional regression network^30^ lies in regressing this Gaussian weighted output matrix from the input image channels. Here, the term “fully-convolutional” refers to the fact that the target variable is a matrix of full image size instead of a being vector quantity. The output matrix yields the center locations of the nuclei upon thresholding locally. The hierarchical (layered) filter structure constitutes a “deep” neural net. The filters encapsulate both linear (convolution) and non-linear (rectified linear unit, ReLU^10^) operations. We adapted an elegant regression framework for cellular object detection^30^. Further, we implemented a simple CNN architecture that minimizes an L2 loss with an exponentially decreasing learning rate^31,32^.

The training process proceeded in two stages^30^, first with a pre-training stage (with cropped 100 ×100 pixel cropped images and augmented by geometric transformations like flipping and rotation), followed by a final training with full images. The test set had five images annotated by six experts. The images were further augmented (by rotation) to make the final test set of 20 images in total.

The deep CNN had layers comprising convolutional kernels and ReLUs. The CNN layers, along with any parameters, were specified in the following order from the input channels to the output^31,32^: convolution (3 ×3 ×2 ×32 kernels), ReLU, convolution (3 ×3 ×32 ×32 kernels), ReLU, convolution (3 ×3 ×32 ×1 kernel). The convolution kernels were initialized with Xavier weights^39^. The peaks of the Gaussian weights in the target matrix were set at 7, following prior convention^30^. The learning rate started at 10-3 for pre-training, and at 10-2 for final training, and decayed exponentially. The weight decay was set at 0.001 for both cases. The scores were thresholded at 1.0, with a window for non-maximum suppression as big as 25 × 25 pixels.

## Acknowledgements

We thank A. Paulson and C. Krishnamurthy for contributing microscopy images analyzed in **Fig.1** and **Fig. 2d**. We thank instructors and students of the 2014 Marine Biology Laboratory Physiology course at Woods Hole for assistance with microtubule gliding assays and image collection (especially R. Fischer). We thank A. Raj and L. Beck for testing Quantius and offering many helpful suggestions. We recognize L. Bugaj, L. Sanders, C. Nilson, and B. Kawas for critical discussion. This work was funded through a Jane Coffin Childs postdoctoral fellowship to A.J.H., the Department of Defense Breast Cancer Research Program (W81XWH-10-1-1023 and W81XWH-13-1-0221 to Z.J.G.), the NIH Common Fund (DP2 HD080351-01 to Z.J.G.), the NSF (MCB-1330864 to Z.J.G.), the Chan-Zuckerberg Biohub Investigator Program (to Z.J.G.), the UCSF Program in Breakthrough Biomedical Research, and the UCSF Center for Cellular Construction (DBI-1548297 to S.K.B., S.B., and Z.J.G.), an NSF Science and Technology Center.

## Author contributions

A.J.H., J.D.M., and Z.J.G. designed Quanti.us. J.D.M. developed and implemented the online website. A.J.H. and D.P.B. analyzed raw Quantius data. S.K.B. and S.B. developed and implemented the machine learning analysis. All authors wrote and edited the manuscript.

